# RalB uncoupled exocyst mediates endothelial Weibel-Palade body exocytosis

**DOI:** 10.1101/2024.09.16.613344

**Authors:** Moua Yang, Alexandra Boye-Doe, Salma A.S. Abosabie, Alexandra M. Barr, Lourdes M. Mendez, Anish V. Sharda

## Abstract

Ras-like (Ral) GTPases play essential regulatory roles in many cellular processes, including exocytosis. Cycling between GDP- and GTP-bound states, Ral GTPases function as molecular switches and regulate effectors, specifically the multi-subunit tethering complex exocyst. Here, we show that Ral isoform RalB controls regulated exocytosis of Weibel-Palade bodies (WPBs), the specialized endothelial secretory granules that store hemostatic protein von Willebrand factor. Remarkably, unlike typical small GTPase-effector interactions, RalB binds exocyst in its GDP-bound state in resting endothelium. Upon endothelial cell stimulation, exocyst is uncoupled from RalB-GTP resulting in WPB tethering and exocytosis. Furthermore, we report that PKC-dependent phosphorylation of the C-terminal hypervariable region (HVR) of RalB modulates its dynamic interaction with exocyst in endothelium. Exocyst preferentially interacts with phosphorylated RalB in resting endothelium. Dephosphorylation of RalB either by endothelial cell stimulation, or PKC inhibition, or expression of nonphosphorylatable mutant at a specific serine residue of RalB HVR, disengages exocyst and augments WPB exocytosis, resembling RalB exocyst-binding site mutant. In summary, it is the uncoupling of exocyst from RalB that mediates endothelial Weibel-Palade body exocytosis. Our data shows that Ral function may be more dynamically regulated by phosphorylation and may confer distinct functionality given high degree of homology and the shared set of effector protein between the two Ral isoforms.

## Introduction

Regulated exocytosis is fundamental to normal functioning of specialized eukaryotic cells.^1^ Examples include blood cells as well as endocrine cells that contain specialized secretory granules, which, in response to specific stimuli, release cargoes that play essential roles in processes such as inflammation and metabolism.^2–4^ Endothelial cells that line the vasculature are critical to maintaining vascular integrity, but also release numerous bioactive products in response to vascular injury by the process of regulated exocytosis. von Willebrand factor (vWF), a large multimeric multifunctional plasma glycoprotein, is one such factor which is indispensable for hemostasis and thrombosis.^5–7^ Deficiency of functional vWF is the most common bleeding disorder encountered in humans, but elevated levels of vWF also increase cardiovascular morbidity.^8,9^ Additionally, vWF plays important roles in angiogenesis, smooth muscle cell proliferation and pathophysiology of vascular inflammation and cancer metastasis.^10–13^ Therefore, regulated localized release of vWF is important for maintenance of vascular integrity but also vital to numerous other pathophysiologic processes.

vWF is stored in specialized secretory granules, Weibel Palade bodies (WPBs).^14,15^ vWF forms C-terminal dimers immediately after synthesis in the endoplasmic reticulum followed by N-terminal multimerization in the *trans-*Golgi network (TGN).^14,15^ Nascent immature WPBs arise from budding at the TGN facilitated by tubulating vWF multimers and then relocate to the periphery of the cell on microtubules, where F-actin anchored WPBs await stimulatory signals to exocytose and release their cargoes extracellularly.^16^ The process of maturation is marked by increasing WPB electron density caused by further multimerization, compaction and organization of the vWF multimers into elongated tubules which impart WPBs their characteristic ‘cigar’ shape, 0.5-5 micron in length.^14^ This process, from biogenesis of WPBs to maturation and ultimately its evoked exocytosis, requires coordination of a large and complex trafficking machinery that remains to be completely defined.^17,18^ Specifically, the mechanisms underlying regulated WPB fusion with the plasma membrane in response to endothelial cell stimulation is not fully understood.^17,18^

Small GTPases, a large family of monomeric proteins that bind and hydrolyze GTP to GDP, act as master regulators of various essential cellular processes including vesicle trafficking.^19^ GDP to GTP exchange and subsequent hydrolysis are accelerated by specific guanine nucleotide exchange factors (GEFs) and GTPase activating proteins (GAPs), respectively. GTP-bound small GTPases bind and activate downstream effectors thereby acting as molecular switches turning on and off corresponding downstream functions. Ral belongs to the Ras GTPase superfamily and exists in two isoforms, RalA and RalB.^20^ Despite a high homology of around 83%, and a shared set of effectors between the two, RalA and RalB possess isoform-specific functions.^21^ This includes actin organization, gene expression, survival, autophagy, endocytosis, and exocytosis.^20^ Consequently, Ral controls many physiologic functions including glucose and lipid metabolism, immune response, neuronal plasticity, oncogenesis, among others.^20,22–25^ Ral has previously been shown to regulate WPB exocytosis.^26,27^ ^28^ Specifically, inhibition of Ral by a small inhibitory peptide resulted in impaired vWF release. However, how Ral regulates WPB exocytosis is not known.

One of the major effectors of Ral in its diverse functions is the exocyst complex.^29–31^ The exocyst complex belongs to a family of multi-subunit tethering complexes that promote granule fusion by physically tethering vesicles to the target membrane.^32^ Exocyst comprises of eight subunits, EXOC1-8 (sec3, sec5, sec6, sec8, sec10, sec15, Exo70 and Exo84 in yeast), and is implicated in vesicular trafficking, exocytosis, cytokinesis, ciliogenesis and cell division.^31^ Ral is not required for exocyst subunits synthesis but regulates exocyst complex assembly on secretory granules and exocyst function.^21,33^ Ral specifically interacts with exocyst subunits EXOC2 and EXOC8 competitively and regulate different cellular functions.^21^ We have previously shown that exocyst complex is present on WPBs and regulates vWF release, but the precise role of exocyst in WPB exocytosis and regulation of its function remains unclear.^34^

This study focused on control of exocyst-dependent WPB exocytosis by Ral GTPase. We find that RalB isoform of Ral GTPase controls WPB exocytosis. GTP-loaded RalB triggers WPB tethering to the plasma membrane and vWF release. Notably, we find that RalB interacts with the exocyst complex in GDP-bound state in resting endothelium and that PKC-dependent RalB phosphorylation modulates interaction with exocyst. Subsequent uncoupling of exocyst from GTP-loaded RalB in activated endothelium triggers WPB tethering to the plasma membrane facilitating WPB exocytosis and vWF release.

## Results

### RalB is required for regulated Weibel-Palade Body exocytosis

We have previously shown that exocyst regulates WPB exocytosis.^34^ Given that exocyst is a major effector of Ral GTPase in its various cellular functions, we first evaluated the effect of Ral depletion on WPB exocytosis.^30^ RalA and RalB can possess distinct functions despite similar enzymology.^25^ Previously, it was reported that Ral regulates WPBs with the use of an inhibitory peptide, but the regulatory role of Ral isoforms in WPB exocytosis requires further characterization.^26–28^ To evaluate this further, we used specific siRNAs to deplete RalA and RalB in HUVECs and studied thrombin-evoked vWF release. We found that depletion of RalB significantly impaired WPB exocytosis (**Figure 1A-B)**. This effect of RalB KD was confirmed using two siRNAs (**Figure S1A-B**). In addition to RNAi, we confirmed the regulatory role of RalB in evoked WPB exocytosis using a small molecule inhibitor of RalB. Dihydroartemisinin, an antimalarial drug, has been previously shown to specifically degrade cellular RalB, but not RalA.^35^ Dihydroartemisinin selectively degraded RalB in HUVECs in a dose dependent manner sparing RalA (**Figure 1C**). This was associated with a significant impairment in thrombin-stimulated WPB exocytosis (**Figure 1D**).

**Figure 1.**
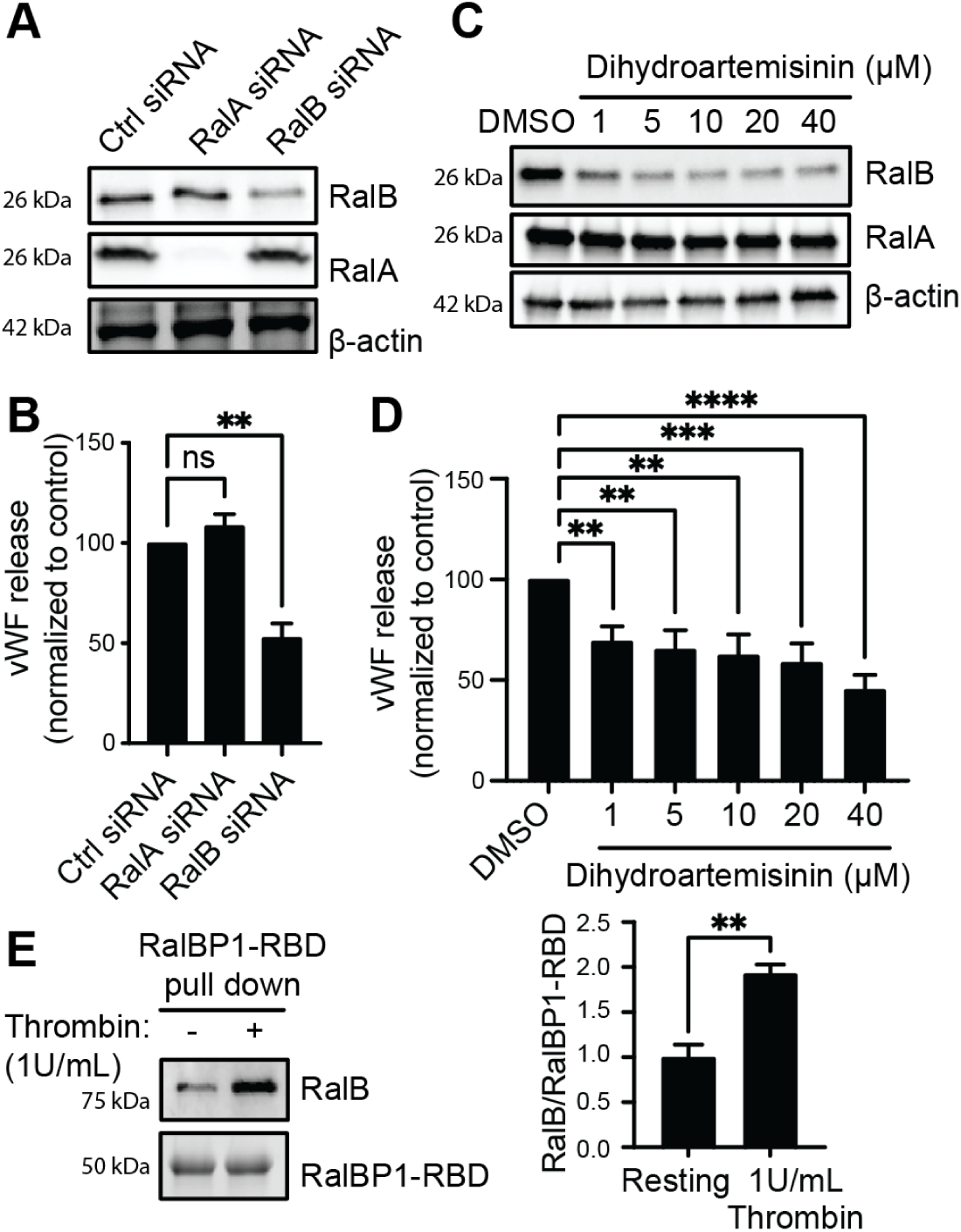
RalB is switched ‘on’ upon thrombin stimulation and regulates endothelial Weibel-Palade body exocytosis. (A) HUVECs depleted of RalA or RalB using specific siRNAs and compared with control siRNA. Western blot of total cell lysates showing RalA and RalB knockdown after 48 hours of transfection of siRNA, with β -actin as loading control. (B) Bar graphs showing total vWF antigen in media as estimated by ELISA 30 minutes after 1 U/ml thrombin stimulation 48 hours after siRNA transfection (\*\**P*<0.01; from 3 individual experiments each with duplicates). (C) HUVECs treated with increasing concentrations of dihydroartemisinin for 2 hours or DMSO. Western blot showing degradation of RalB but not RalA by dihydroartemisinin, with β-actin as loading control. (D) Bar graphs showing total vWF antigen in media as estimated by ELISA 30 minutes after 1 U/ml thrombin stimulation 2 hours after dihydroartemisinin treatment (\*\**P*<0.01, \*\*\**P*<0.001; \*\*\*\**P*<0.0001 from 3 individual experiments each with duplicates). (E) Total cell lysates of resting and thrombin-treated (1 U/ml for 2 minutes) HUVECs were incubated with RalBP1-RBD conjugated agarose slurry and eluates immunoblotted with anti-RalB antibody to determine total GTP-loaded RalB in each condition. Coomassie stain showing RalBP1-RBD as loading control (left). Bar graphs showing densitometric analysis (right). (\*\**P*<0.01; from 3 individual experiments).

Next, we wanted to determine the activation status of RalB, or in other words GDP/GTP cycling, in resting versus stimulated HUVECs. The activation of Ral in regulated exocytosis is believed to be Ras-independent.^20^ Secretory cell stimulation results in release of RalGDS, a Ral GEF, from beta-arrestin and consequently RalGDS catalyzes GTP exchange in Ral thereby switching Ral ‘on’.^20,28^ Using a functional assay that depends on binding of RalB-GTP to the Ral binding domain (RBD) of its specific effector Ral binding protein 1 (RalBP1), we estimated the amount of RalB-GTP in resting versus thrombin-stimulated HUVECs. RalB binds RalBP1-RBD only in GTP-loaded state.^36,37^ Our data shows that RalB is GDP-loaded in resting HUVECs, but GTP-loaded upon thrombin stimulation (**Figure 1E**). This implies that endothelial stimulation switches ‘on’ RalB which subsequently facilitates WPB exocytosis. 2-minute time point was chosen based on previous literature showing maximal Ral activation within this time frame.^38^

### RalB associates with mature WPBs

Next, we wanted to determine the localization of RalB with respect to WPBs. Ral has been previously reported to be present on WPBs.^27^ To determine if this is RalB, we performed confocal immunofluorescence microscopy (IF) with anti-RalB and anti-vWF antibodies on fixed permeabilized HUVECs. WPBs appear as peripherally distributed cigar-shaped structures. RalB was found to colocalize with vWF on WPBs (**Figure 2A**). RalB also colocalized with other markers of mature WPBs, specifically P-selectin and Angiopoietin-2 (**Figure S2**).

**Figure 2.**
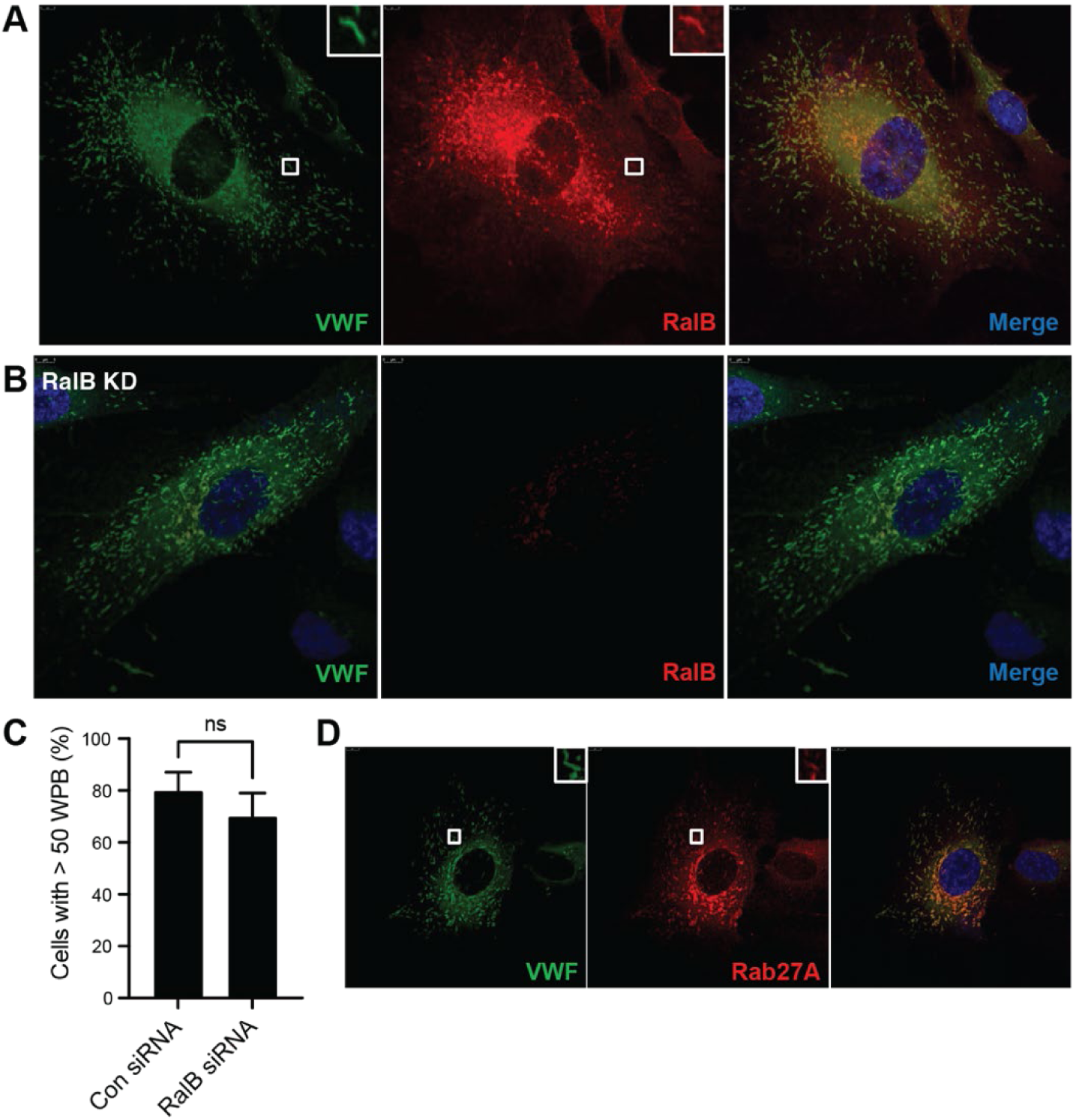
RalB associates with mature WPBs. (A-C) IF performed on HUVECs with anti-vWF and anti-RalB antibodies 48 hours after transfection with RalB siRNA compared with control siRNA. (A) Micrograph showing presence of RalB (red) in WPBs labeled with vWF (green) in control HUVECs. DAPI (blue) showing nuclei. Inset depicting magnified image of a single WPB. Scale bar 5 μm. (B) Micrograph showing unaffected distribution of WPBs (vWF; green) and RalB (red) in RalB-depleted HUVECs. Scale bar 5 μm. (C) Bar graph showing number of WPBs per cell 48 hours after transfection with RalB siRNA and compared with controls (ns=not significant; 30 random cells examined for each condition in 3 individual experiments). (D) IF performed with anti-vWF and anti-Rab27a antibodies 48 hours after transfection with RalB siRNA. Micrograph showing presence of Rab27A (red) in WPBs labeled with vWF (green). DAPI (blue) showing nuclei. Inset depicting magnified image of a WPB. Scale bar 5 μm.

Depletion of many components of WPB trafficking machinery can impair WPB maturation.^34^ To see if RalB may also play a role in WPB maturation, we performed IF in RalB-depleted HUVECs with anti-vWF antibody. RalB depletion did not affect localization or number of WPBs as compared with control siRNA (**Figure 2B-C**). Mature WPB cigars in RalB depleted cells showed typical peripheral distribution and were positive for Rab27a, a marker of WPB maturity (**Figure 2D**). This implies that RalB does not participate in WPB biogenesis and is likely recruited to mature WPBs.

### Expression of autoactivated RalB targets WPBs to the plasma membrane and augments vWF release

After small GTPase activation, GTPase activating proteins (GAPs) bind GTP-loaded small GTPases and increase the intrinsic rate of GTP hydrolysis.^20^ RalGAP is the GAP specific to Ral GTPases and binds Ral rapidly following Ral GTP-loading.^20^ The glycine to valine mutation at the 23^rd^ residue of Ral (Ral-G23V) eliminates RalGAP binding to both Ral isoforms.^24^ Consequently, Ral-G23V exists primarily in GTP-bound state and is therefore autoactivated. To determine the effect of expression of autoactivated RalB on WPB exocytosis, we expressed RalB-G23V under the control of a CMV promoter using lentiviral transduction in HUVECs. RalB-G23V was generated by site-directed mutagenesis of pLA-N-Flag-RalB-WT plasmid. RalBP1-RBD pulldown assay, as described above, confirmed GTP loading of RalB-G23V (**Figure S3).** HUVECs expressing RalB-G23V demonstrated significant augmentation of vWF release as compared with HUVECs expressing RalB-WT even in the absence of thrombin stimulation (**Figure 3A**).

**Figure 3.**
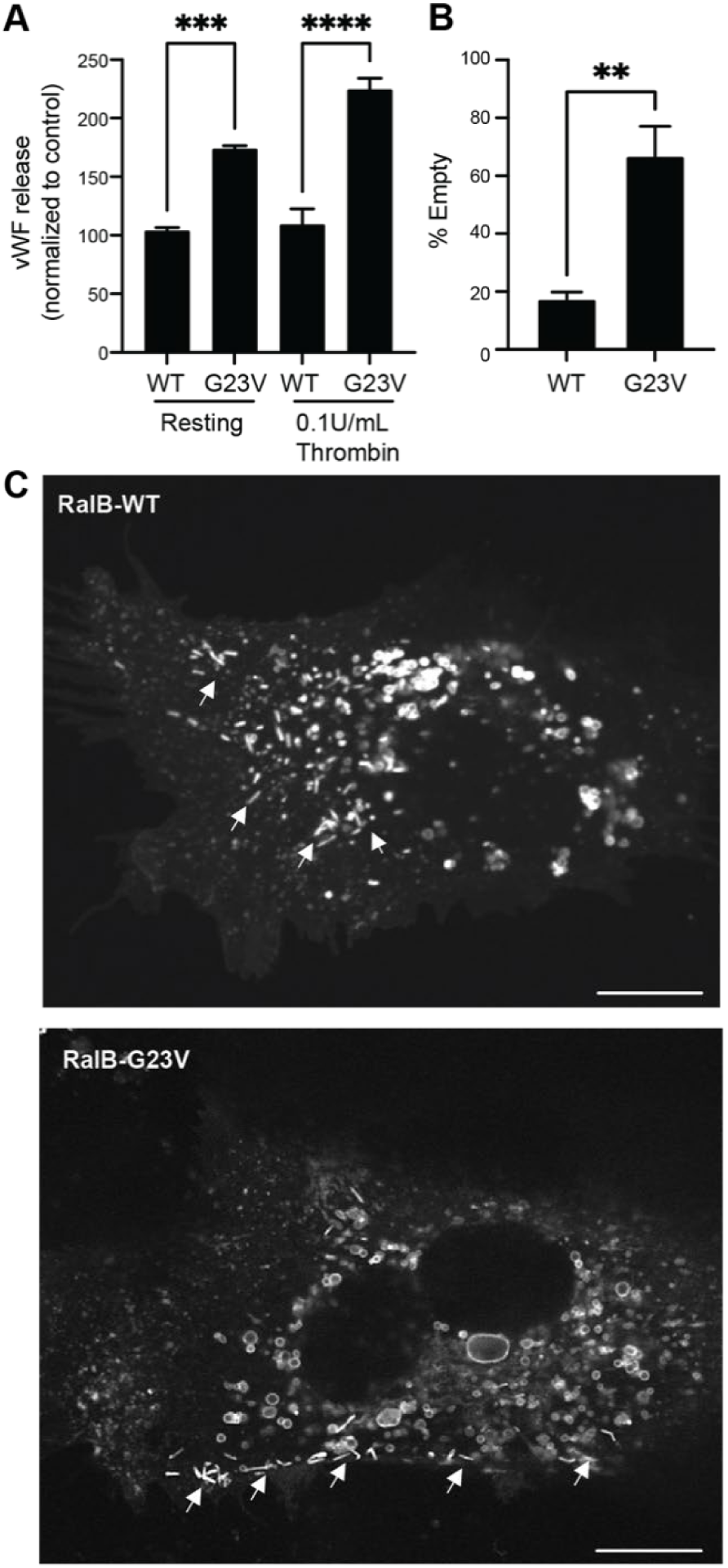
RalB activation tethers WPBs to the plasma membrane. (A) Expression of autoactivated RalB (RalB-G23V) in HUVECs augments vWF release as compared with RalB-WT. Bar graphs showing total vWF antigen in media estimated by ELISA from resting and thrombin stimulated HUVECs (1 U/ml thrombin for 30 minutes) (\*\*\**P*<0.001 or \*\*\*\**P*<0.0001, from 3 individual experiments each with triplicates). (B) RalB-G23V and RalB-WT expressing HUVECs labeled with anti-vWF antibody and imaged with a confocal immunofluorescence microscope. 100 random cells counted, and cells expressing < 15 WPBs considered as empty cell. Bar graphs showing percent empty cells (\*\**P*<0.01; from 3 individual experiments). (C) CD63-EGFP expressing HUVECs transduced with lentiviral particles containing RalB-G23V or RalB-WT. WPBs appear as elongated ‘cigars’ (white arrowheads) distributed among round CD63-positive endolysosomes. WPBs appear tethered to the plasma membrane in RalB-G23V cells as compared with cytoplasmic distribution in RalB-WT cells. Scale bar 10 μm.

Next, we expressed autoactivated RalB-G23V in HUVECs expressing CD63-EGFP and performed live cell microscopy. CD63 is present on mature WPBs and we have previously described the use of CD63-EGFP expressing HUVECs for this purpose.^34^ We expressed RalB-WT and RalB-G23V under the control of a CMV promoter using lentiviral transduction in CD63-EGFP HUVECs. RalB-G23V cells had higher number of ‘empty cells’, defined as <15 WPB per cell as compared with RalB-WT cells (**Figure 3B**).

Notably, WPBs in RalB-G23V cells were found at the plasma membrane, as compared with cytoplasmic distribution in RalB-WT HUVECs (**Figure 3C**). Many WPBs in RalB-G23V expressing HUVECs congregated at the plasma membrane in groups of two to three, reminiscent of cumulative WPB fusion.^17^ Live cell microscopy showed typical random motility in RalB-WT expressing HUVECs as previously described, whereas WPBs in RalB-G23V expressing cells were found to be relatively immobile, tethered at the plasma membrane (**Supplementary videos 1 and 2**).^39^ These data suggest that RalB directly regulates WPB tethering and fusion.

### EXOC2 associates with RalB-GDP in resting endothelium

Given exocyst is a major Ral effector, We next determined if RalB-dependent WPB tethering noted above was being mediated by exocyst. First, we performed microscopy to localize RalB and exocyst on WPBs. Confocal IF confirmed presence of both RalB and exocyst on WPBs in paraformaldehyde fixed and permeabilized HUVECs (**Figure 4A**). Additionally, we performed immune-electron microscopy (EM) to study the localization of RalB and exocyst ultrastructurally. Unfortunately, the specific monoclonal anti-RalB antibody demonstrated poor labeling with glutaraldehyde fixation used for immunogold labeling.

**Figure 4.**
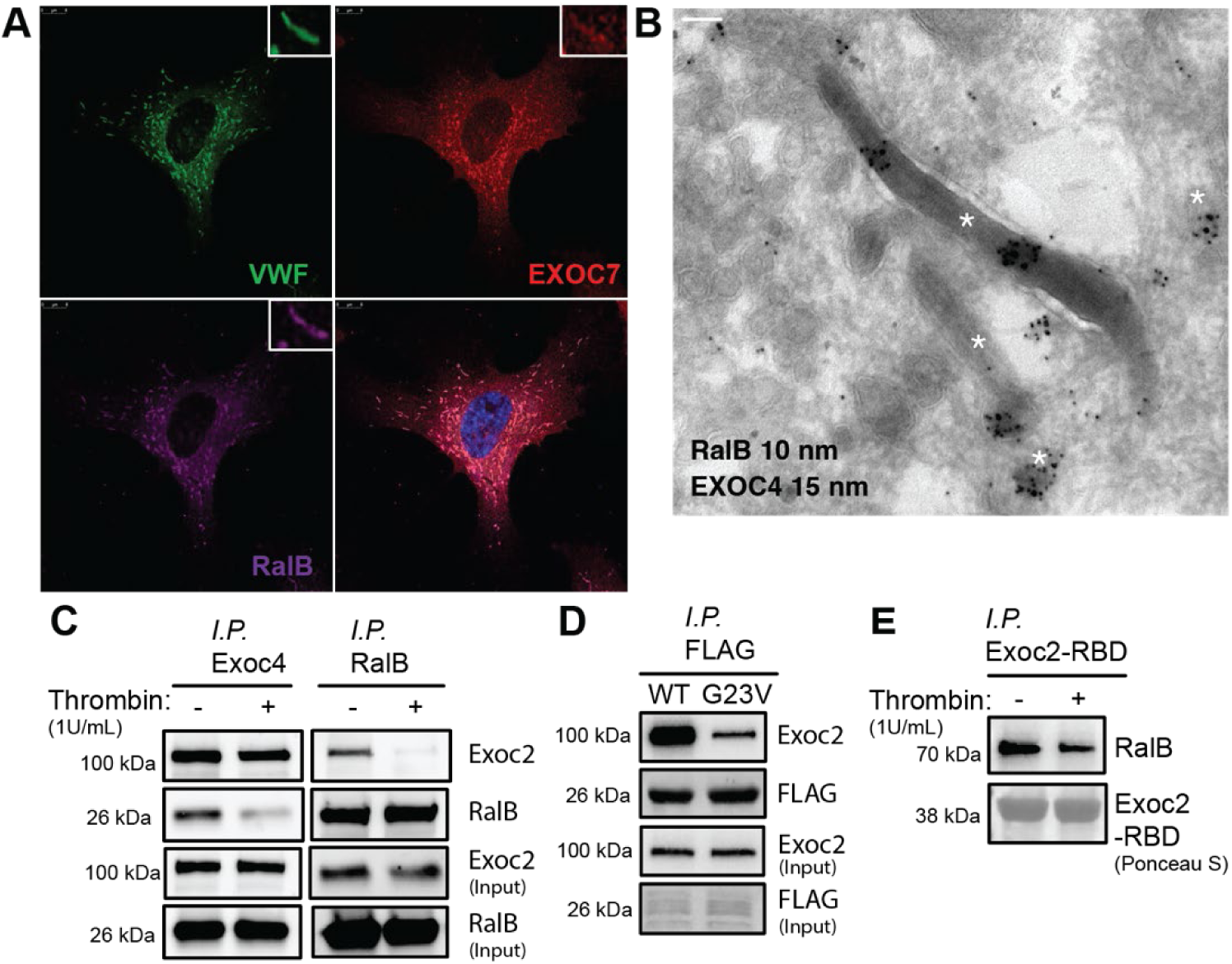
Exocyst interacts with GDP-loaded RalB. (A) Confocal immunofluorescence micrographs showing presence of RalB (purple) and EXOC7 (red) with vWF (green) in mature WPBs. Insets depict magnified image of a mature WPB. DAPI (blue) showing nucleus. Scale bar 8 μm. (B) Transmission electron microscopy of HUVECs with double immunogold labeling with anti-EXOC4 (15 nm) and FLAG (10 nm) antibodies. Transmission electron micrograph showing both EXOC4 and RalB decorating mature electron-dense WPB ‘cigars’ (white asterisk). Scale bar 100 nm. (C) Western blots showing immunoprecipitation (IP) of resting and thrombin stimulated (1U/ml for 2 min) total HUVEC lysates with anti-RalB or anti-EXOC4 antibodies resolved by SDS-PAGE and immunoblotted with anti-RalB and anti-EXOC2 antibodies. Total HUVEC lysate (∼5% of IP) used for input. (D) Total cell lysates of RalB-WT and RalB-G23V expressing HUVEC incubated with magnetic beads conjugated with anti-FLAG antibody and eluates immunoblotted with anti-FLAG and anti-EXOC2 antibodies. (E) Total cell lysates of resting and thrombin-treated (1 U/ml for 2 minutes) HUVECs incubated with EXOC2-RBD conjugated agarose slurry and eluates immunoblotted for with anti-RalB antibody. Ponceau S staining showing EXOC2-RBD as loading control.

Therefore, we used HUVECs expressing RalB-WT, as above, which possesses an N-terminal DYKDDDDK (FLAG) tag. Double immunogold labeling with anti-FLAG and anti-EXOC4 antibodies followed by transmission EM confirmed the presence of RalB and exocyst on mature WPB cigars (**Figure 4B**).

To determine RalB-exocyst interaction biochemically, we first performed immunoprecipitation experiments in native HUVEC lysates. Based on the generally accepted paradigm of the interaction of small-GTPases with their effectors in GTP-loaded, or switched ‘on’ state, we carried out immunoprecipitation studies in both resting and thrombin-stimulated HUVEC lysates. RalB was immunoprecipitated in HUVEC lysates using anti-RalB antibody bound dynabeads, and then eluates resolved by SDS-PAGE and immunoblotted for RalB and EXOC2. Immunoprecipitation confirmed RalB interaction with EXOC2, but quite unexpectedly, RalB interacted with EXOC2 in resting HUVECs where RalB is predominantly GDP-bound (**Figure 4C**). This dynamic interaction was confirmed by reverse immunoprecipitation studies. Again, RalB interacted with exocyst in resting HUVEC lysates (**Figure 4C**). Anti-EXOC4 antibody performed better than anti-EXOC2 antibody for this assay. Since exocyst is an obligate multisubunit complex, this does not influence our findings.

To corroborate this in cells expressing autoactivated RalB (RalB-G23V), we performed FLAG pulldown in RalB-WT and RalB-G23V HUVEC lysates. Using anti-FLAG antibody-conjugated magnetic beads, RalB-WT and RalB-G23V were immunoprecipitated, eluates resolved by SDS-PAGE and then immunoblotted for RalB and EXOC2. Like the native HUVEC lysates, EXOC2 precipitated out significantly more with RalB-WT as compared with RalB-G23V (**Figure 4D**). We also sent eluates from FLAG pulldown experiments for mass spectrometry analysis. EXOC2 was identified as a RalB interacting protein, but the peptide counts were significantly higher in RalB-WT as compared with RalB-G23V eluates (**Supplementary file 1**). In addition to EXOC2, exocyst subunit EXOC8 is also known to bind RalB, but we did not identify EXOC8 in our immunoprecipitates.^20^

### Ral binding domain of EXOC2 preferentially binds RalB-GDP

Given the data presented above, which show that EXOC2 preferentially interacts with GDP-loaded RalB in total HUVEC lysates, we wanted to confirm these results by studying interaction of RalB with recombinant EXOC2 Ral binding domain (EXOC2-RBD). The N-terminal 99 residues of EXOC2 form its RBD.^40,41^ The original interaction between EXOC2-RBD and Ral was determined with RalA, showing that EXOC2-RBD preferentially bound GTP-loaded RalA in solution. Notably, RalA was truncated to remove its C-terminal hypervariable region (HVR) in these previous reports. It has since become clear that Ral HVR, which spans residues 177-202 imparts critical functional differences between RalA and RalB, and modulates Ral effector interaction.^20^

We cloned EXOC2-RBD into a bacterial expression vector containing GST tag and expressed it in *E. coli* (**Figure S4A)**. Pulldown studies on resting and thrombin-stimulated HUVEC lysates were performed with purified EXOC2-RBD GST, and eluates immunoblotted for RalB. Like the data above, EXOC2-RBD precipitated significantly more RalB in resting compared with thrombin-stimulated HUVEC lysates (**Figure 4E**). This confirms that exocyst preferentially binds GDP-loaded RalB. Treatment of lysates with GDP or non-hydrolysable GTP analogue GTPγS did not affect this interaction of EXOC2 with RalB-GDP in resting endothelial lysates (**Figure S4B**). This suggests that other regulatory factors, such as RalB post-translational modifications, likely modulate RalB-exocyst interaction.

### PKC-dependent RalB phosphorylation modulates exocyst binding

Post-translational modifications of small GTPases are known to modulate effector binding and regulation.^42,43^ Post-translational modifications of Ral, such as phosphorylation and ubiquitination, are also known to regulate GDP/GTP-binding and effector binding.^44–46^ PKC dependent phosphorylation of serine residues at positions 192 and more so 198 in RalB HVR were previously shown to affect RalB-GTP loading and RalBP1 and exocyst binding in a colorectal cancer cell line in resting conditions.^46^ Therefore, we wanted to determine if RalB phosphorylation could be modulating the unexpected binding of EXOC2 with RalB-GDP in regulated WPB exocytosis we discovered in both resting and stimulated endothelium. Thrombin is known to activate multiple PKC isoforms and phosphatases in endothelial cells.^47–49^ To explore this further, we studied the effect of pharmacologic PKC modulation on RalB phosphorylation and exocyst binding. HUVECs were treated with bryostatain-1, a PKC activator, as well as sotrastaurin, a pan-PKC inhibitor, lysed, and immunoprecipitated with anti-RalB antibodies. The precipitates were resolved by SDS-PAGE and blotted with anti-phosphoserine, anti-RalB or anti-EXOC2 antibodies. Our data shows that at basal state in resting HUVECs, RalB is phosphorylated, and thrombin stimulation dephosphorylates RalB (**Figure 5A**). EXOC2 preferentially bound phosphorylated RalB in resting endothelium, as previously observed. Treatment with PKC inhibitor sotrastaurin reduced RalB phosphorylation, reducing EXOC2 binding in resting cells. Conversely, PKC activator bryostatin-1 increased RalB phosphorylation, enhancing exocyst binding even in thrombin-stimulated cells (**Figure 5A**).

**Figure 5.**
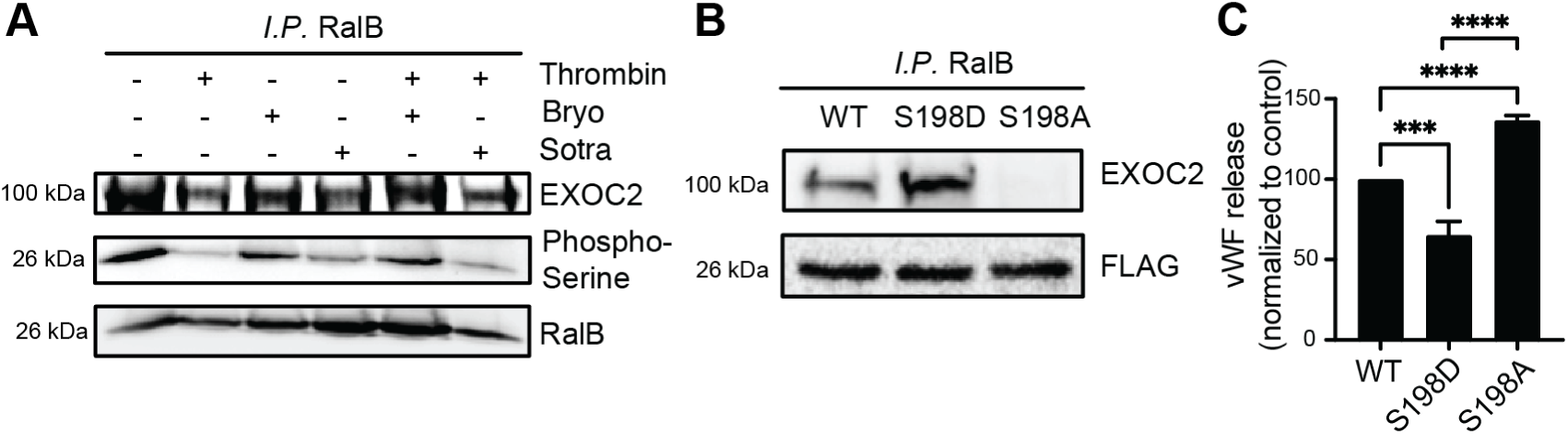
PKC-dependent RalB phosphorylation modulates exocyst binding and WPB exocytosis. (A) Thrombin stimulation dephosphorylates RalB and reduces EXOC2 binding. IP with anti-RalB or anti-EXOC4 antibodies of total cell lysates of resting and thrombin stimulated HUVECs (1U/ml for 2 min) with and without PKC activator bryostatin-1 (100nM for 10 min) or PKC inhibitor sotrostaurin (500nM for 1 hour), resolved by SDS-PAGE and immunoblotted with anti-phosphoserine, anti-RalB and anti-EXOC2 antibodies. Total HUVEC lysate (∼5% of IP) used as input. (B) Expression of phosphomimetic RalB-S198D mutant increases EXOC2 binding, and phosphodeficient RalB-S198A mutant decreases EXOC2 binding in HUVECs. Total cell lysates of RalB-WT, RalB-S198D and RalB-S198A expressing cells incubated with magnetic beads conjugated with anti-FLAG antibody and eluates immunoblotted with anti-Flag and anti-EXOC2 antibodies. (C) Bar graphs showing total vWF antigen in media (at 30 minutes) collected from resting HUVECs expressing RalB-WT, RalB-S198D and RalB-S198A as estimated by ELISA (\*\*\**P*<0.001; \*\*\*\**P*<0.0001, from 3 individual experiments each with triplicates).

We confirmed the effect of post-translational phosphorylation of RalB on EXOC2 binding and WPB exocytosis by expressing RalB with serine to alanine (non-phosphorylatable) and serine to aspartic acid (phosphomimetic) mutations at position 198. S198, and to a lesser extent S192, phospho-mutations located in RalB HVR were previously shown to modulate RalB function.^46^ Employing site-directed mutagenesis, we generated S198D and S198A mutations in RalB-WT, and transduced HUVECs with lentiviral particles carrying these constructs. Compared with RalB-WT, RalB-S198D enhanced exocyst binding, while RalB-S198A decreased exocyst binding (**Figure 5B**). The effect of these mutations at S198 on WPB exocytosis reflected the status of RalB-exocyst binding. RalB-S198D mutation resulted in an impairment in WPB exocytosis, whereas RalB-S198A mutation significantly augmented WPB exocytosis and vWF release (**Figure 5C**). Combined, these data suggest that EXOC2 is a substrate of phosphorylated RalB and that PKC-dependent post-translational phosphorylation of RalB modulates its interaction with EXOC2 affecting downstream cellular functions.

### Uncoupling exocyst from RalB triggers WPB tethering and vWF release

Our interpretation of these data is that it is the uncoupling of exocyst from GTP-loaded RalB in stimulated endothelial cells that triggers WPB tethering and exocytosis. To evaluate this further, we generated aspartate to glutamate mutation at position 49 of RalB (RalB-D49E), which has been previously shown to eliminate Ral-exocyst interaction.^36,50^ Employing site-directed mutagenesis, we generated D49E mutation in both RalB-WT and RalB-G23V, and transduced HUVECs with lentiviral particles carrying these constructs. As expected, RalB-D49E mutation impaired EXOC2 binding (**Figure 6A**). RalB-G23V expressing HUVECs had low baseline RalB-exocyst interaction, as already noted above, which was eliminated with addition of D49E mutation. Next, we determined vWF release in these cells by estimating vWF antigen levels in culture media. Consistent with our hypothesis, D49E mutation significantly augmented basal vWF release as compared with RalB-WT (**Figure 6B**). Addition of D49E mutation did not further augment vWF release in RalB-G23V cells that already had low RalB-exocyst binding.

**Figure 6.**
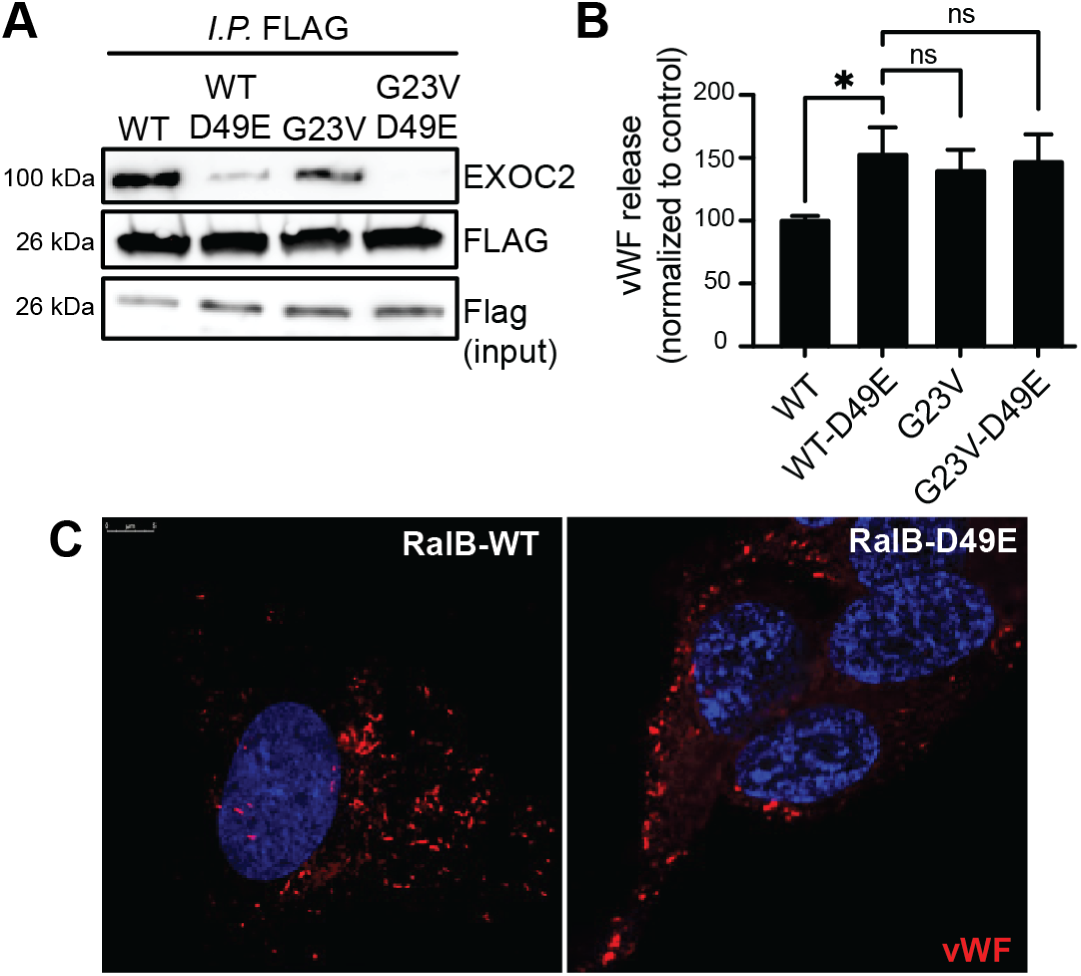
RalB uncoupled exocyst triggers WPB exocytosis. (A) Expression of RalB exocyst binding site mutation in HUVECs (RalB-WT D49E and RalB-G23V D49E). Total cell lysates of RalB-WT and RalB-G23V and their respective D49E mutants incubated with magnetic beads conjugated with anti-FLAG antibody and eluates immunoblotted with anti-FLAG and EXOC2 antibodies. (B) Bar graphs showing total vWF antigen in media from HUVECs expressing RalB-WT, RalB-WT D49E, RalB-G23V or RalB-G23V D49E mutants (\**P*<0.01; ns=non-significant; from 3 individual experiments each with triplicates). (C) RalB-WT and RalB-WT D49E expressing HUVECs labeled with anti-vWF antibody and imaged with a confocal immunofluorescence microscope. Micrograph showing WPBs labeled with vWF (red) decorating plasma membrane in RalB-WT D49E mutants as compared to cytoplasmic distribution in RalB-WT. DAPI (blue) depicting nuclei. Scale bar 5 μm.

Lastly, we performed IF in RalB-D49E expressing HUVECs to locate WPBs. RalB-WT and RalB-WT-D49E cells were grown on coverslips, fixed and permeabilized, and labeled with anti-vWF antibody. As compared to the normal cytoplasmic distribution of mature WPBs in RalB-WT cells, WPBs in the presence of RalB-WT-D49E mutation were found decorating the plasma membrane, as previously noted in RalB-G23V expressing HUVECs (**Figure 6C**). These data confirm that RalB uncoupled exocyst triggers WPB tethering and facilitates exocytosis.

## Discussion

Acting as molecular switches, Ral small GTPases integrate extracellular stimuli with downstream cellular processes, including vesicular trafficking, through specific effectors. In this work, we have made two interdependent but equally important observations. First, we provide evidence that Ral isoform RalB positively regulates WPB exocytosis directly through its effector exocyst complex. But more notably, we demonstrate that exocyst can interact with GDP-bound RalB, as in resting endothelium. Once switched ‘on’, exocyst disengages from GTP-bound RalB and consequently RalB uncoupled exocyst triggers WPB tethering to the plasma membrane facilitating WPB exocytosis and vWF release. Moreover, this dynamic association of RalB with exocyst and regulation of its tethering function is modulated by post-translational phosphorylation of RalB (**Figure 7**).

**Figure 7.**
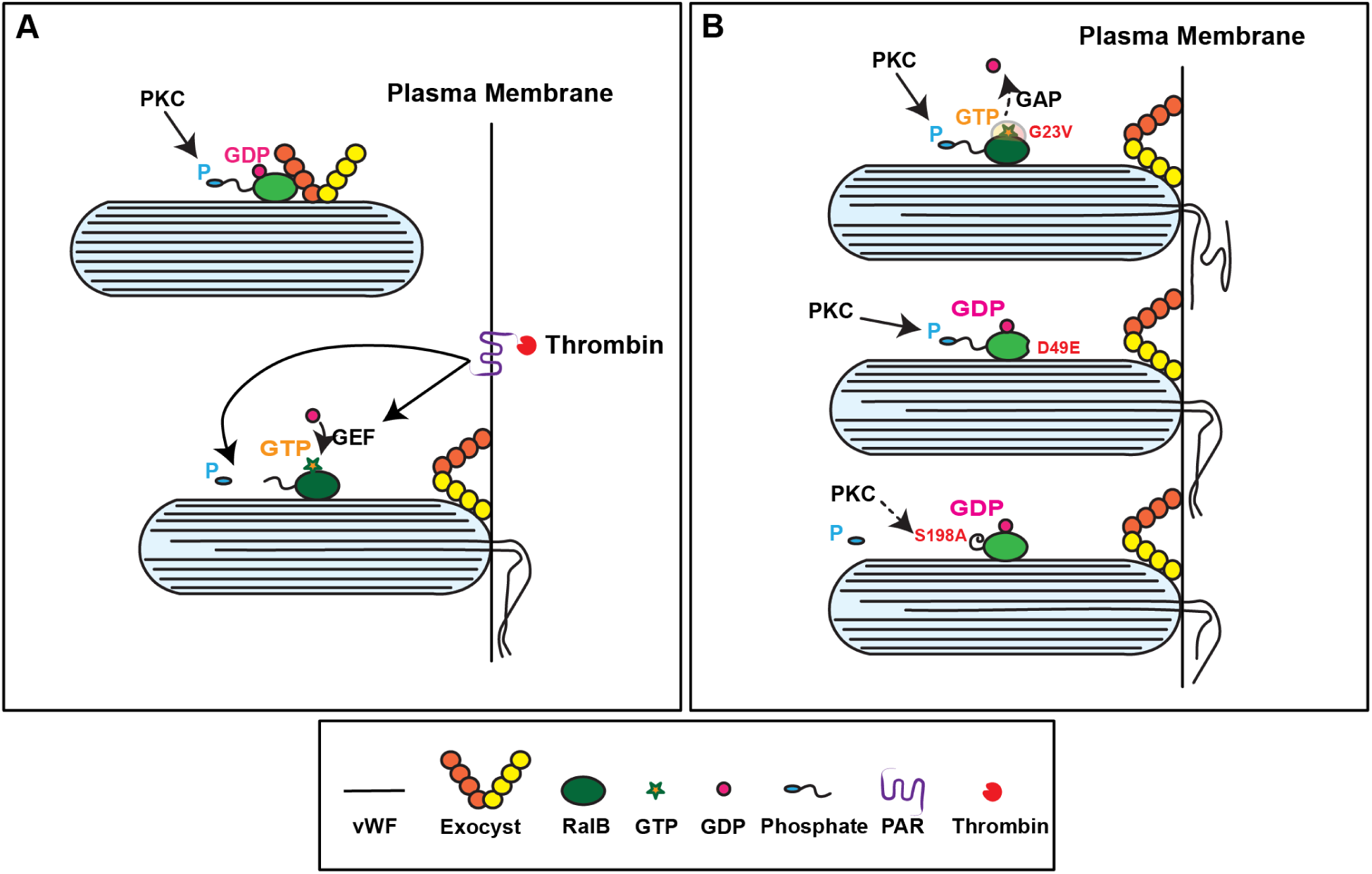
Proposed model for RalB-exocyst dependent WPB tethering. (A) In resting HUVECs, RalB is GDP-loaded as well as phosphorylated by protein kinase C in its C-terminal hypervariable region. Upon endothelial cell stimulation by thrombin, RalB becomes GTP-loaded catalyzed by GEF, as well as dephosphorylated by activation of phosphatases. Consequently, exocyst is uncoupled from dephosphorylated and GTP-bound RalB and tethers WPBs to the plasma membrane facilitating WPB exocytosis and vWF release. (B) RalB mutations which render RalB autoactivated (G23V; top), phosphodeficient (S198A; middle) or uncoupled from exocyst (D49E; bottom), trigger WPB tethering and vWF release in the absence of endothelial cells stimulation. (GEF, guanine nucleotide exchange factor; GAP, GTPase activating protein; P, phosphate; PKC, protein kinase C; PAR, protease activated receptor)

The paradigm for small GTPases as molecular switches is their dual-mode conformation - the GTP-loaded ‘on’ state, where small GTPases bind their effectors, triggering downstream functions, and GDP-loaded ‘off’ state. Our data supports a role for activated GTP-bound RalB triggering endothelial WPB exocytosis, but contrary to the accepted paradigm, by disengaging with its effector exocyst. There are other sporadic examples of GDP-loaded GTPase binding its effector. For example, Slp4a preferentially interacts with GDP-bound Rab27a to inhibit dense-core vesicle exocytosis in PC12 cells.^51^ The effector binding occurs at the switch domains that is conserved in all small GTPases including Ral.^20^ The conformational change that occurs with GTP/GDP cycling acts as a sensor and allows for specific interaction with effectors, typically with GTP-bound forms of GTPases.^52^ The binding site of EXOC2 is also located in the switch I region of Ral, and previously, RalA was reported to bind and control exocyst function in insulin secretion as well as GLUT4 trafficking when GTP-loaded.^20,53,54^ In contrast to these studies, our data clearly shows that exocyst interacts with GDP-bound RalB in resting endothelial cells and disengages from GTP-bound RalB following endothelial cell stimulation and intracellular calcium mobilization. We confirmed this by showing interaction of native RalB with purified EXOC2 Ral binding domain preferentially from resting lysates as compared to stimulated lysates. Furthermore, treatment with either GDP or non-hydrolysable GTP analogue GTPγS did not alter this dynamic interaction.

There is increasing understanding of how posttranslational modifications in small GTPases modulate both GDP/GTP cycling as well as effector binding. Post-translational modifications, particularly phosphorylation, regulates effector selection and affinity for many known Rabs but also Ral.^20,42,43^ Phosphorylation has been shown to regulate both RalA and RalB activity in cell and function specific manner, in addition to ubiquitination in the case of RalB.^45,46,55^ Our data also shows that PKC-dependent phosphorylation of RalB modulates exocyst binding in HUVECs. Specifically, modifications of S198 in RalB HVR affects exocyst binding and WPB exocytosis. The phosphodeficient S198A mutation reduces exocyst binding augmenting WPB exocytosis, and on the other hand phosphomimetic S198D mutation enhances exocyst binding impairing WPB exocytosis. Structural determinants of Ral-EXOC2 interaction have also been described previously.^40,41^ These studies showed that Ral binds exocyst when GTP-loaded, but notably, these studies used a truncated version of RalA with its C-terminal HVR removed. This would eliminate the phosphorylation sites in both RalA and RalB that are so critical in regulating Ral function allosterically.

In addition to Ral phosphorylation modulating exocyst interaction, post-translational modifications of exocyst have also been shown to affect its interaction with RalA, which we have not evaluated in this study.^56^ Both EXOC2 and EXOC8 bind RalB and may do so competitively, however, EXOC8 was not identified in our pulldown data.^45^ The functional significance of this remains unclear. Ral has been previously shown to be required for assembly of exocyst on secretory granules.^57^ Given that RalB depletion impairs endothelial WPB exocytosis, it is likely that RalB is required for exocyst assembly on WPBs, but the precise mechanism remains unclear. Also uncertain is the interplay of other small GTPases that are present on mature WPBs, particularly Rab27a that is also known to bind and regulate exocyst.^51,58–61^ Lastly, the role of RalBP1, a Ral effector that acts as a GAP for Rho GTPases which regulate actin dynamics, remains to be determined in WPB exocytosis.^20^

Pharmacologic modulation of endogenous vWF release would be beneficial in treatment of von Willebrand disease but also in other pathophysiologic conditions where vWF plays essential roles, but the molecular regulation of regulated WPB secretion remains elusive. We propose that RalB uncoupled exocyst mediates endothelial WPB exocytosis and vWF release, and that Ral phosphorylation is a major determinant of Ral-exocyst interaction. Given high degree of homology and the shared set of effectors between RalA and RalB, post-translational modifications may confer distinct functionality to these GTPases.

## Materials and Methods

### Reagents

Fatty acid-free bovine serum albumin, Tween-20, 2-mercaptoethanol, triton X-100, methanol, hydrochloric acid, 0.1% gelatin solution and Bryostatin-1 were obtained from Millipore Sigma. DynaBeads Protein G, MaxiSorp immuno plate, 1-Step^TM^ Ultra TMB-ELISA, 4’,6-diamidino-2-phenylindole dihydrochloride (DAPI), Restore^TM^ Western Blot Stripping Buffer, puromycin dihydrochloride, bacitracin, and lipofectamine RNAiMax as well as 3000 were obtained from Invitrogen. Laemmli sample buffer, 4-15% TGX SDS-polyacrylamide gels were obtained from BioRad. EBM-2 basal medium, human umbilical vein endothelial cells (HUVECs), and HUVECs bullet kits were obtained from Lonza. Human α-thrombin was obtained from Haematologic Technologies. Blotto non-fat dry milk was obtained from ChemCruz. Calcium- and Magnesium-free phosphate buffered saline (PBS), pH 7.4, Opti-MEM reduced serum media, DMEM, fetal bovine serum and penicillin/streptomycin were obtained from Gibco. Streptavidin-HRP was obtained from Abcam. Sotrastaurin was obtained from Selleckchem.

### Antibodies

The primary antibodies are listed in the table below.

| Antigen | Host | Vendor |
| --- | --- | --- |
| RalA | Rabbit monoclonal | Abcam (ab12667) |
| RalB | Rabbit monoclonal | Abcam (ab129077) |
| EXOC2 | Rabbit polyclonal | Proteintech (12751-1-AP) |
| EXOC4 | Mouse monoclonal | MilliporeSigma (MABC570) |
| EXOC7 | Mouse monoclonal | Kerafast (ED2001) |
| vWF | Rabbit polyclonal | Dako |
| vWF | Mouse monoclonal | Thermo Fisher (MA5-14029) |
| FLAG | Mouse monoclonal | SigmaMillipore (F1804) |
| P-selectin | Mouse monoclonal | Thermo Fisher (14-0628-82) |
| Phosphoserine | Rabbit polyclonal | SigmaMillipore (AB1603) |
| β-Actin | Mouse monoclonal | Thermo Fisher (MA1-140) |

Species specific HRP-conjugated secondary antibodies as well as Verablot secondary for immunoprecipitation experiments was obtained from Abcam. Species specific Alexa Fluor 488, 568 and 647 antibodies were obtained from Invitrogen.

### Cell Culture

Human umbilical vein endothelial cells were cultured in EBM-2 basal media supplemented with bullet kit containing endothelial growth supplements (complete media) in gelatin-coated tissue-culture treated flasks or plates in 37C incubator with 5% CO2. Only passage 1-5 were used for experiments.

### Plasmids, RNAi, transfection, and transduction

RalA (s11758) and RalB (s11761 and s11762) siRNA were purchased from Life Technology. For siRNA-transfection, HUVECs were grown to confluency on 6 or 12-well plates in complete media. siRNAs were prepared by mixing 10 µM of negative control, RalB, or RalA siRNA with lipofectamine RNAiMax transfection reagent in OptiMEM low-serum media for 10 min to a final concentration of 5-10 pM, prior to the addition to the cells. The transfection media was replaced with complete media after 6-8 h of transfection. Experiments were performed on RalB and RalA knockdown cells 48 h post-transfection. pLA-CMV-N-Flag-RalBWT (plasmid#50990), pLX301 (plasmid#25895), pEGFP-C2-CD63 (plasmid#62964) were obtained from Addgene.

To make lentiviruses, first, 12 µg of plasmid was incubated with 25 µl of lipofectamine 3000 together with packaging plasmids (9 µg psPAX2 (addgene#12260), 3 µg PMD2 (addgene#12259), 3 µg pRSV-Rev (addgene#12253)) in 1 ml of Opti-MEM low serum medium for 45 min. Lenti-X-293T cells cultured to 70-80% confluency in 10 cm dishes in 9 ml of growth media were then transfected with this mix.

Growth media was replaced after overnight transfection. Lentivirus was then harvested at 48 h and 72 h post-transfection and either added directly to HUVECs in culture or concentrated with Lenti-X concentrator (Takara) and stored in -70C for future use.

For viral transduction, P1 HUVECs were grown to 80-90%% confluency and either the growth media was replaced with freshly harvested virus-containing media or with frozen concentrated virus in complete media. After overnight incubation, media was replaced with fresh complete media. Antibiotic selection was performed after 48 hours of infection.

### vWF ELISA

HUVECs were grown on 6- or 12-well tissue culture plates until confluent. The cells were serum starved for 90 min at 37°C prior to stimulation with 1U/mL human alpha thrombin for 30 min. The media containing releasates were immediately frozen or further processed for vWF antigen detection. A 96-well flat bottom MaxiSorp immuno plate was coated with 1 µg/mL anti-vWF antibody (Dako rabbit polyclonal) in a sodium carbonate/bicarbonate buffer pH 9.5 for 1 h at room temperature prior to blocking with 3% w/v BSA in PBS-T for 30 min. Pooled plasma from healthy human donors containing vWF antigen were used as a standard by serial dilution in PBS. VWF antigen in the diluted releasates were then captured onto the wells for 1 h at room temperature. The antibody-vWF complexes were washed with PBS-T (0.1%) three times followed by incubation with an anti-vWF antibody (Dako rabbit polyclonal) conjugated with biotin for 60 min at room temperature. 100 ng/mL streptavidin-HRP was added in 1% w/v BSA in PBS-T for 45 min at room temperature followed by five washes with PBS-T. TMB solution was then added until noticeable development of color. The reaction was stopped with 1N hydrochloric acid and spectrophotometrically read at 450 nm.

### Site-directed mutagenesis and TOPO cloning

Site-directed mutagenesis was carried out either using Q5 site-directed mutagenesis kit (New England Bio) or In-fusion cloning (Takara) according to the manufacturers’ protocols. Primers for site-directed mutagenesis were designed based on the protocol employed.

pEGFP-C2-CD63 was TOPO cloned using gateway cloning into pLX301 vector using pENTR/D-TOPO cloning kit (Life Technology) and primers were designed based per kit protocol. RalB-binding domain (RBD) of EXOC2 was also TOPO cloned into pET16 vector using pENTR/D-TOPO cloning kit.

### Confocal microscopy

Confocal immunofluorescence microscopy was performed on HUVECs cultured on chambered coverslips, fixed with 4% PFA for 10 min, permeabilized with 0.25% triton X-100 for 10 min, blocked with 3% w/v BSA, incubated with primary antibodies overnight at 4°C, followed by fluorophore-conjugated secondary antibodies for 45 min and counterstained with DAPI for 2 min. Coverslips were mounted using Prolong Gold mounting media and imaged with a LEICA SP8 upright microscope with 63X 1.35 normal aperture oil objective. LAS-X software was used to analyze imaging data.

Live-cell imaging was carried out on cells plated on glass chambered coverslips with a Yokagawa spinning disk confocal on an inverted Nikon Ti fluorescence microscope fitted with OkoLab 37°C, 5%CO_2_ cage microscope incubator with 100X 1.25 normal aperture oil objective. Images were further processed in ImageJ.

### Electron Microscopy

Immunogold-electron microscopy was performed based on the Tokayasu method. Briefly, for preparation of cryosections, the cells were removed from the dish with Trypsin, fixed with 4% paraformaldehyde + 0.1% glutaraldehyde (in 0.1M Sodium Phosphate buffer, pH 7.4), washed with PBS and then placed in PBS containing 0.2M glycine to quench free aldehyde groups. (15min). Samples were then frozen in liquid nitrogen and sectioned at -120°C and the sections were transferred to formvar/carbon coated copper grids. The double immunogold labeling was carried out at room temperature on a piece of parafilm. All antibodies and protein A gold (University Medical Center, Utrecht, the Netherlands) were diluted in 1% BSA in PBS. Grids were blocked with 1% BSA for 10 minutes and incubated with primary antibodies for 30 minutes. Double immunogold labeling was completed by sequential incubation with the first species-specific secondary antibody followed by 15nm Protein-A gold, a fixation step in between, and finally the other secondary antibody followed by10 nm Protein-A gold. The grids were examined in a JEOL 1200EX electron microscope and images were recorded with an AMT 2k CCD camera.

### Immunoprecipitation

Lysates for immunoprecipitation (IP) were prepared using IP lysis buffer (Invitrogen) or magnesium lysis buffer (Millipore Sigma). Lysates were incubated with 2.5 µg anti-RalB or EXOC4 antibody-bound protein G Dynabeads or Anti-Flag M2 magnetic beads overnight, washed with PBS-T three times and boiled with reduced Laemmli sample buffer. Eluted immunoprecipitates were either sent for mass spectrometry or resolved with SDS PAGE followed by immunoblotting.

Immunogold-electron microscopy was performed based on the Tokayasu method as previously described.^34^

## Acknowledgements

The authors thank Dr. Mary Munson and Dr. Michael Marks for their guidance throughout this work. This work was supported by the National Institute of Health [grant numbers K08HL150246 to A.V.S., R00HL164888 to M.Y.], and the American Society of Hematology Scholar Award [to M.Y.].

## Author Contributions

M.Y. designed and performed experiments, analyzed data, and edited the manuscript. A.B., S.A. and A.M.B. performed experiments. L.M.M designed experiments and edited the manuscript. A.V.S. designed and performed experiments, analyzed data, and wrote the manuscript.

## Conflict of Interest

The authors declare no financial conflict of interest.

---

**Figure S1.**
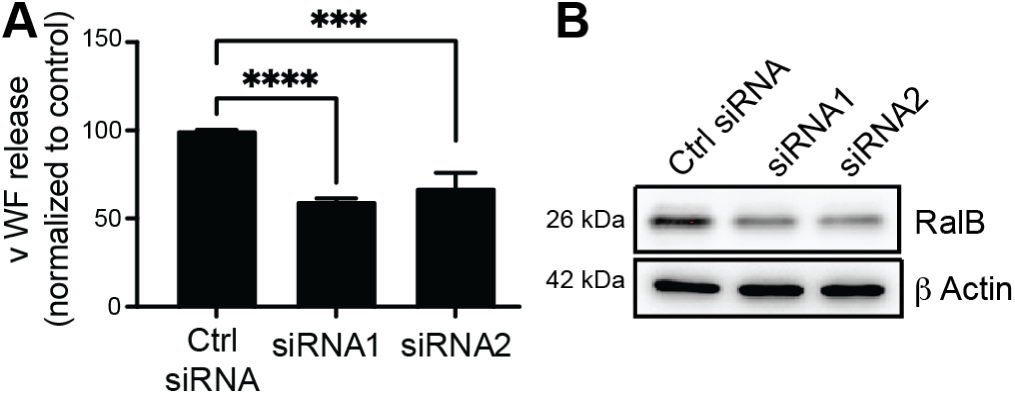
RalB regulates WPB exocytosis. (A) HUVECs depleted of RalB using two specific siRNAs and compared with control siRNA. Bar graphs showing total vWF antigen in media 30 minutes after 1 U/ml thrombin stimulation 48 hours after siRNA transfection as estimated by ELISA (\*\*\**P*<0.001; \*\*\*\**P*<0.0001; 3 individual experiments each with duplicates). (B) Western blot of total cell lysates showing RalB knockdown at 48 hours, with β-actin as loading control.

**Figure S2.**
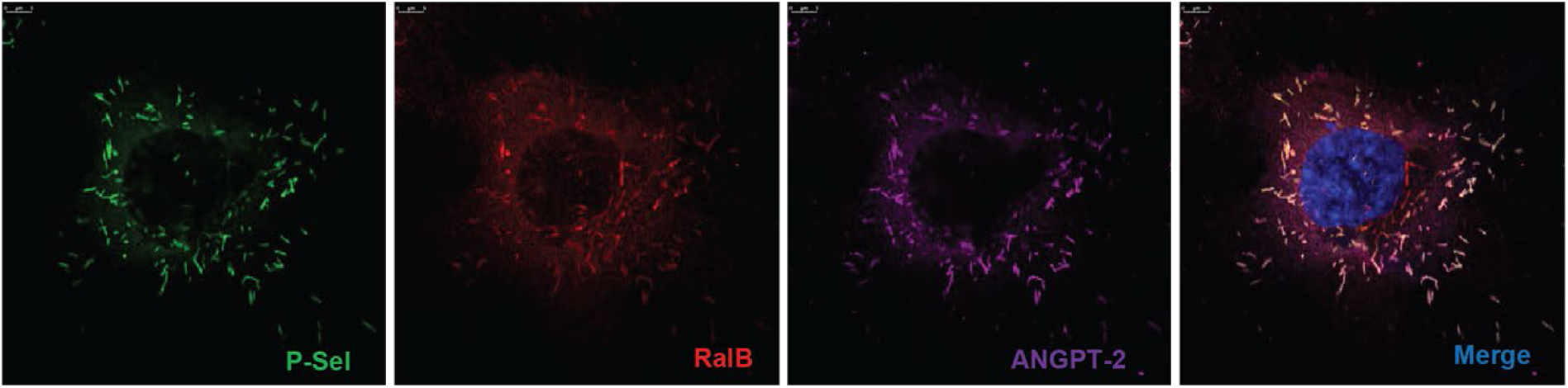
RalB associates with mature WPB. (A) Confocal immunofluorescence micrographs showing presence of RalB (red) in WPBs labeled with P-selectin (green) and Angiopoietin-2 (purple). DAPI (blue) showing nuclei. Scale bar 5 μm.

**Figure S3.**
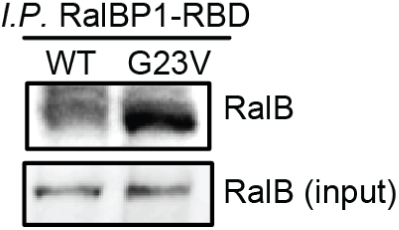
RalB-G23V is GTP-bound. Total cell lysates of HUVECs expressing RalB-WT and RalB-G23V incubated with RalBP1-RBD conjugated agarose slurry and eluates immunoblotted with anti-RalB antibody to quantify GTP-loaded RalB. ∼5% of lysates used for input.

**Figure S4.**
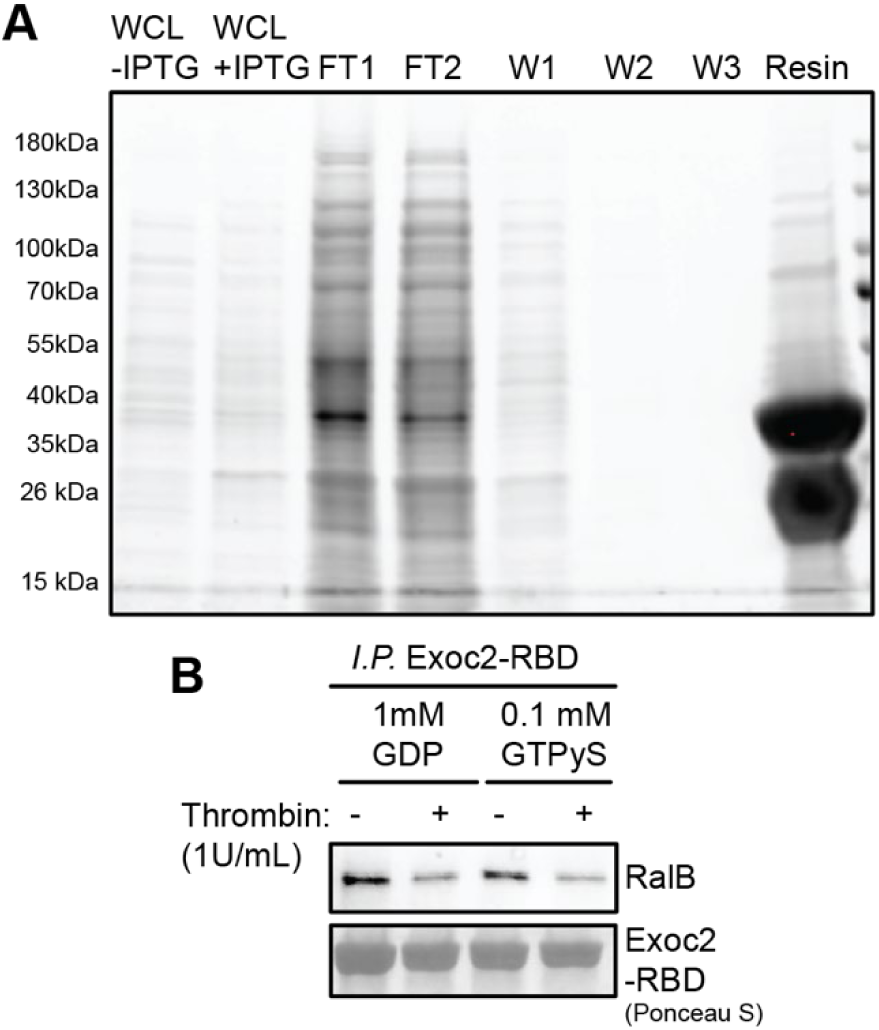
(A) SDS-PAGE gel followed by Coomassie staining to show purification of recombinant GST-tagged EXOC2-RBD in the whole cell lysate, flow through, washes, and bound to the resin. WCL, Whole Cell Lysate; IPTG, Isopropyl β-d-1-thiogalactopyranoside; FT1, Flow Through 1; FT2, Flow Through 2; W1, Wash 1; W2, Wash 2; W3, Wash 3. (B) Total cell lysates of resting and thrombin-treated (1 U/ml for 2 minutes) HUVECs treated with either GDP (1mM) or GTPγS (100μM) for 30 minutes, followed by incubation with EXOC2-RBD conjugated agarose slurry at 4°C for 30 minutes. Eluates immunoblotted with anti-RalB antibody. Ponceau S staining sowing EXOC2-RBD as loading control.

**Supplementary Videos 1 and 2.** CD63-EGFP expressing HUVECs were transduced with RalB-WT (Video 1) or RalB-G23V (Video 2) and analyzed by live cell spinning disk confocal microscopy. Images acquired at 1 frame per second. WPBs are evident in both conditions, in addition to CD63-EGFP positive endosomes. While WPBs appear distributed in the cytoplasm and show typical back-and-forth random motility in RalB-WT HUVECs, WPBs appear tethered to the plasma membrane and relatively immobile in RalB-G23V HUVECs.

## Notes

### Competing Interest Statement

The authors have declared no competing interest.

